# Identification and Evaluation of a Panel of Strong Constitutive Promoters in *Listeria monocytogenes* for Improving the Expression of Foreign Antigens

**DOI:** 10.1101/2021.02.09.430548

**Authors:** Junfei Ma, Qianyu Ji, Shuying Wang, Jingxuan Qiu, Qing Liu

## Abstract

Attenuated *Listeria monocytogenes* (*L. monocytogenes*) could be used as a vaccine vector for immunotherapy of tumors or pathogens. However, the lack of reliable promoters limits its ability to express foreign antigens. In this work, 21 promoters from *L. monocytogenes* were identified by RNA-seq analysis under two conditions of pH 7.4 and pH 5.5. Based on the constructed fluorescence report system, 7 constitutive promoters showed higher strength than that of P_help_, a previously reported strong promoter. Further, the selected 5 constitutive promoters also showed high activity in the production of UreB, a widely used antigen against *Helicobacter pylori* (*H. pylori*). In particular, a well-characterized constitutive promoter P_18_, which performed best in both fluorescence intensity and UreB production, was proved to be highly active in vitro and in vivo. In summary, we provide a useful promoter library for *Listeria* species and offer a reference for constitutive promoter mining in other organisms.

**Key points:**

- 21 promoters from *L. monocytogenes* were identified by RNA-seq.
- Fluorescent tracer of *L. monocytogenes* (P_18_) was performed in vitro and in vivo.
- A well-characterized constitutive promoter P_18_ could improve the expression level of a foreign antigen UreB in *L. monocytogenes*

## Introduction

*L. monocytogenes* is a gram-positive bacterium that could stimulate innate response and cellular immune responses including CD4^+^ T cells and CD8^+^ T cells[1, 2]. Due to the endogenous pathway of antigen processing, attenuated *L. monocytogenes* such as *L. monocytogenes*Δ*actA/inlB* strain was widely applied as a vaccine vector in the research of microbial immunotherapy for controlling tumor and infectious diseases[3–6]. To date, in the construction of vaccine strain of attenuated *L. monocytogenes*, the promoter of *hly* gene (P_*hly*_) was generally used to elicit the expression of foreign antigens[7, 8]. In the previous study, we found that P_*hly*_ was not strong enough to meet the expression levels of the foreign antigen and further provoke a specific immune response[9]. P_help_, modified from P_*hly*_, has higher promoter strength, but it is still to be improved[10]. Besides, the live vaccine based on *L. monocytogenes* vector is forced to bear the acidic environments like the macrophage phagosome and tumor microenvironment[11, 12]; the oral *L. monocytogenes*-vector vaccine also encounters stomach acid environment[13]. This requires the promoters to be stable enough to adapt to the acidic environments. Thus, one of the key limitations is the lack of a panel of well-characterized promoters to regulate the expression of foreign antigen in *L. monocytogenes*.

Constitutive promoters are widely used for fine-tuning the expression levels of key genes in metabolic engineering and synthetic biology[14, 15]. Usually, constitutive promoters are screened from the promoters of essential genes because of their constant transcript levels[16]. Hence, a method based on RNA-seq analysis could be applied to obtain native promoters according to different transcriptional strength. More recently, in the field of synthetic biology, this method has been developed to construct the native promoter libraries of various industrial microorganisms such as *Streptomyces albus, Streptococcus thermophilus,* and *Pseudomonas*[17–19]. A panel of strong constitutive promoters could be obtained by selecting the potential promoters based on RNA-seq analysis and evaluating their activities based on fluorescent reporter genes.

With the increasing interest in *L. monocytogenes* in microbiological immunotherapy of tumors and pathogens, the development of transcriptome sequencing technology offers an opportunity to mine reliable constitutive promoters. In this work, a panel of constitutive promoters was identified based on the systematic analysis of transcriptome data of *L. monocytogenes* cultivated in different pH conditions. The green fluorescent protein (GFP) reporter was used for identifying the characterization of these promoters, and the activity of a well-characterized promoter was evaluated in vitro and in vivo. Based on our further needs, several promoters with different strengths were selected for experimentally evaluating the production of UreB, a widely used antigen against *H. pylori*. These constitutive promoters enriched the promoter library of *Listeria*, which should be of great value for metabolic engineering and synthetic biology in this genus.

## 2 Materials and methods

### 2.1 Strains, plasmids, and medium

All bacteria strains and plasmids used in this study were listed in Table S1. *Escherichia coli* DH5α was grown on Luria-Bertani broth (LB) medium at 37 ◻. It was applied for plasmid construction and propagation. *L. monocytogenes* wild-type EGD-e and EGD-eΔ*actA/inlB* were cultured in brain heart infusion medium (BHI, Beijing Land bridge, China) at 37 °C.

### 2.2 Growths of *L. monocytogenes* under different pH conditions

The overnight-grown wild-type EGD-e strain was collected by centrifugation at 5000 × g at 4 ◻, washed in PBS (10 mM, pH 7.4), and adjusted to 1.0 at OD_600_ nm. The cultures were then diluted 1:100 in fresh BHI broth (pre-adjusted to pH 4.5, 5.0, 5.5, 6.0 or 7.4, respectively) to 0.2 at OD_600_ nm (3.6×10^8^ CFU/mL). The growth curve was assessed for 12 h at 37 ◻ with shaking. The bacterial solution was taken for determination at OD_600_ nm and 1-h interval using a SpectraMax M2 microplate reader (Molecular Devices, USA). Three independent experiments were performed and the results were reported as average.

### 2.3 Acid resistance determination of pre-acid treated *L. monocytogenes*

Based on the growths of wild-type EGD-e under different pH conditions, eight kinds of pre-acid stress treatments (pH 4.5, 30 min; pH 4.5,1 h; pH 5.0, 30 min; pH 5.0, 1 h; pH 5.5, 3 h; pH 5.5, 6 h; pH 6.0, 3 h; and pH 6.0, 6 h) were performed to explore acid resistance of EGD-e for screening the optimal treatment. Due to that the bacteria was barely growing at pH 4.5 and 5.0, the initial inoculum at pH 4.5 and 5.0 was 1.0×10^9^ CFU/mL (OD_600_ = 0.4), while that at pH 5.5 and 6.0 was 3.6×10^8^ CFU/mL (OD_600_ = 0.2). The acid-treated cultures were collected by centrifugation at 5000 × g for 10 min at 4 ◻, washed in PBS, and adjusted to 0.4 at OD_600_ nm. The cultures were then plated onto BHI agar after appropriate dilutions for counting surviving bacteria. At the same time, 1 mL of cultures (OD_600_=0.4) were harvested, washed once in PBS, resuspended in isometric BHI broth (pre-adjusted to pH 3.0, death acidity), and cultured at 37 ◻ for 20 min. After that, the cultures were collected, resuspended in isometric BHI broth (pH 7.4), and plated onto BHI agar. The acid resistance of bacteria was characterized by survival rate, which was defined as the ratio of survival number after acid lethal treatment to that after acid lethal treatment. Survival rates are reported as the mean of three independent experiments, each performed in duplicate.

### 2.4 RNA-seq

The wild-type EGD-e strain was cultured in BHI medium under two conditions of pH 7.4 and 5.5 at 37 ◻ for 3 h with shaking. The initial inoculum was 3.6×10^8^ CFU/mL. Cultures were collected by centrifugation at 5000 × g at 4 ◻, and washed in DEPC water. Total RNA was extracted using Trizol reagent (Invitrogen, USA) and then ribosomal RNA was removed by a Ribo-Zero Magnetic kit (Epicentre Biotechnologies, USA). RNA quality was checked by NanoDrop (ThermoFisher Scientific, USA) followed by RNA degradation and contamination verification on 1% agarose gel. The cDNA libraries were prepared using the TruSeq RNA Library Preparation kit (Illumina, USA) and sequenced on Illumina Hiseq 4000 platform by Sangon Biotechnology Co. Ltd., Shanghai, China. Clustering and sequencing were performed by Sangon that employed spliced reads to determine connectivity. FPKM (Fragments Per Kilobase of gene model per Million mapped reads) was calculated using featureCounts. FPKM, which also considers the effect of sequencing depth and gene length on reads counts, is a commonly used method for evaluating gene expression levels[20]. Promoter strength could be reflected by FPKM.

### 2.5 Construction of plasmids and strains

The green fluorescent protein (GFP) gene was amplified from plasmid pUC57-GFP and the promoters of 25 highly expressed genes were cloned from wild-type EGD-e genome. Every cloned promoter and amplified *gfp* sequence were inserted into plasmid pERL3 using ClonExpress® MultiS One Step Cloning Kit (Vazyme, Nanjing, China) to create a series of pERL3 derivatives, pERL3-P_1_-GFP to pERL3-P_25_-GFP. widely used promoters P_*hly*_ and P_help_, as controls, were applied to generate pERL3-P_*hly*_-GFP and pERL3-P_help_-GFP using the same method. All primers used in this study were listed in Table S2. The promoter sequences were listed in Table S3. The constructed plasmids with different promoters were proliferated in *E. coli* DH5α, identified by sequencing, and transformed into wild-type EGD-e using electroporation.

### 2.6 Measurement of fluorescence intensity of GFP

The constructed strains were cultured for 12 h, 24 h, 36 h, and 48 h, respectively. Cultures were collected by centrifugation, washed twice with 20mM Tris-HCl, and suspended. After adjusting to the appropriate absorption at 600 nm (OD_600_), the fluorescence intensity of GFP (excitation at 485 nm and emission at 525 nm) was measured in microtiter plates (Assay Plate, 96 wells, Black Polystyrene; Corning, USA) using a SpectraMax M2 microplate reader (Molecular Devices, USA).

### 2.7 Fluorescent tracer of *L. monocytogenes* in macrophage RAW264.7

EGD-e and EGD-eΔ*actA/inlB* carrying the plasmid pERL3-P_18_-GFP were chosen for fluorescent tracer of *L. monocytogenes* in macrophage RAW264.7. Approximately 2×10^5^ RAW264.7 cells were seeded on cover glass (Thermo Fisher Scientific, USA) in a 12-well plate per well overnight. The cells were infected with bacteria at the multiplicity of infection (MOI) of 100 for 2 h. After washing three times, gentamicin was added for 30 min to eliminate the extracellular bacteria. Then cells were fixed with 4% paraformaldehyde in PBS at room temperature for 30 min, and permeabilized in 0.1% TritonX-100 in PBS for 5 min. Actin-stain 488 (Red, Cytoskeleton Inc.) and DAPI (Blue, H-1200, Vector Lab.) were utilized to stain actin and label the cell nucleus, respectively. Images were observed and captured on Leica DM 2500 fluorescence microscope (Leica, Germany).

### 2.8 Fluorescent tracer of *L. monocytogenes* in vivo

In vivo imaging of EGD-eΔ*actA/inlB* carrying fluorescent reporting plasmid was tested in traditional C57BL/6 mice. P_18_, as the strongest promoter, was selected to observe its activity in vivo. EGD-e carrying the plasmid pERL3-P_18_-GFP were cultured overnight, washed, and resuspended using PBS. Mice were inoculated with EGD-eΔ*actA/inlB* (5×10^7^ CFU/mouse) by intravenous injection. Three days postinoculation, the mice were anesthetized with isoflurane and imaged in a Perkin Elmer IVIS Lumina II system.

### 2.9 Measurement of Urease B subunit (UreB) production

The promoters with different strengths P_18_, P_7_, P_12_, P_9_, P_24_, P_help_, and P_*hly*_ were chosen for evaluating their abilities to express foreign antigen UreB. The plasmids pERL3-P_18_/P_7_/P_12_/P_9_/P_24_/P_help_/P_*hly*_-UreB were constructed and transformed into EGD-eΔ*actA/inlB.* The constructed strains were cultured overnight and the total soluble proteins were extracted by ultrasonic crushing apparatus. The expression levels of UreB under different promoters in EGD-eΔ*actA/inlB* were evaluated by western blotting probed with a mouse anti-UreB polyclonal antibody. At the same time, the purified UreB proteins with gradient dilution as standard were set for quantitative analysis of UreB in EGD-eΔ*actA/inlB* by gray scan using ImageJ. In this part, the quantitation of UreB was based on total protein normalization and the proteinic concentration was assayed by BCA Kit (Solarbio, China).

### 2.10 Statistical analyses

All data were analyzed with the GraphPad Prism 5 software (GraphPad Software Inc., La Jolla, CA, USA) and expressed as mean ± standard deviation (SD). Statistical significance was tested using one-way ANOVA. P < ․05 was considered as statistically significant (*P < ․05, **P < ․01, ***P < ․001, ****P < ․0001; ns: not significant).

## 3 Results

### 3.1 Growths of *L. monocytogenes* under different pH conditions

Acid stress could delay or stop the growth of *L. monocytogenes*. As shown in Fig. 1 a, compared with the normal condition (pH 7.4), the growth rate of wild-type EGD-e was inhibited by acid stress. The growth at pH 6.0 was slightly inhibited that its stationary phase was delayed by 4 hours and the bacterial concentration of stable period was about 0.68 at OD_600_ nm, which was still close to that at pH 7.4 (0.71). The growth at pH 5.5 was moderately inhibited. The growths at pH 5.0 and 4.5 were severely inhibited, which were barely changed.

**Fig. 1.**
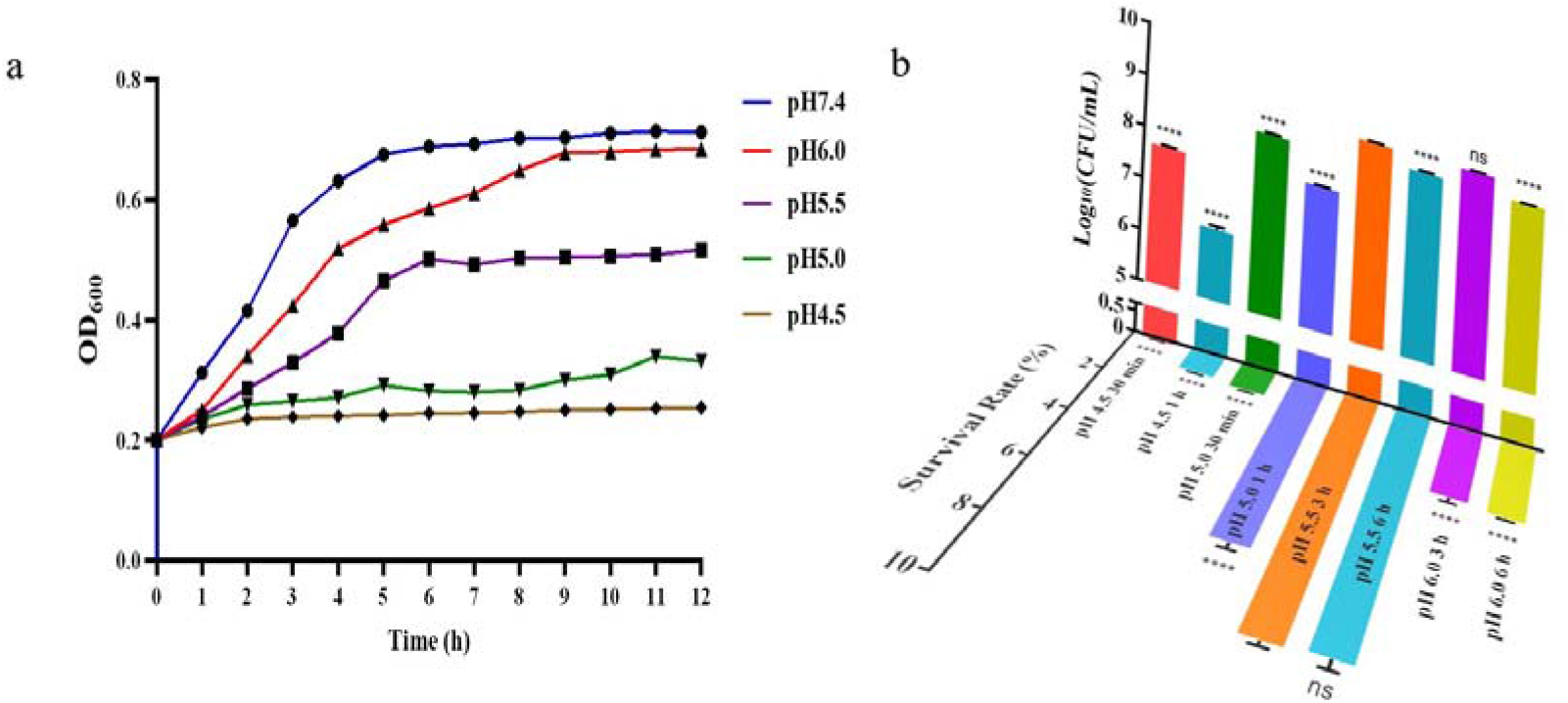
The screening of acid stress treatments for *L. monocytogenes* RNA-seq analysis. **a**. The growth characteristics of *L. monocytogenes* at different pH conditions. The initial inoculation amount of *L. monocytogenes* was 3.6×10^8^ CFU/mL. **b**. Acid resistance determination of *L. monocytogenes* with different acid stress treatments. The viable bacteria number under eight kinds of different acid stress (pH 4.5, 30 min; pH 4.5,1 h; pH 5.0, 30 min; pH 5.0, 1 h; pH 5.5, 3 h; pH 5.5, 6 h; pH 6.0, 3 h; and pH 6.0, 6 h) and survival rate after further acid lethal treatment (pH 3.0, 20min) were measured. Bacteria were adjusted to OD_600_ nm of 0.4 for counting. The error bars indicate the standard deviations from three independent replicates. Statistical significance was compared to the group of pH 5.5, 3 h: ns, no significant, ****, P<0.0001.

### 3.2 The optimal acid stress treatments for *L. monocytogenes* RNA-seq analysis

Acid-inducing tolerance response under sublethal conditions could improve the survival rate of *L. monocytogenes* under acid lethal conditions[21, 22]. In this study, the survival rate was to characterize bacterial acid resistance. The bacteria with stronger acid resistance have higher expression levels of acid stress-associating proteins. As shown in Figure 1 b, the surviving bacteria after pre-acid treatments (pH 5.5, 3 h; pH 6.0 3 h) were 9.13×10^8^ and 9.33×10^8^ CFU/mL respectively, which were close to that under the normal condition (1.0×10^9^ CFU/mL). The lethality of bacteria after these two acid stress pretreatments was extremely low. The surviving bacteria were reduced to 4.70 × 10^8^ and 4.43 × 10^8^ CFU/mL respectively, when the acid induction (pH 5.5 and 6.0) time was extended to 6 h. The surviving bacteria of the other acid stress treatments (pH 4.5, 30 min; pH 4.5,1 h; pH 5.0, 30 min; pH 5.0, 1 h) were 4.9×10^7^, 2.5×10^6^, 3.33×10^8^, and 6.6×10^7^ CFU/mL respectively, which indicated that the strong-acid treatments have higher lethality even for a shorter time.

Further, after acid lethal treatment, the survival rates of pre-acid treatments (pH 5.5, 3 h; pH 5.5, 6 h) were 8.98% and 9.01%, which were the top 2 of all pre-acid treatments. And there was no significant difference between 3 h and 6 h at pH 5.5. A similar trend could be seen at pH 6.0 that the survival rates of pre-acid treatments (pH 6.0, 3 h; pH6.0, 6 h) were both about 3.0%. Besides, the survival rates of pre-acid treatments (pH 5.0, 1 h) could reach 6.8%, while those of the other pre-acid treatments (pH 4.5, 30 min; pH 4.5,1 h; and pH 5.0, 30 min) were 0.12%, 0.97%, and 1.22%. As a whole, the pre-acid treatment (pH 5.5, 3 h), which did well in both bacterial activity and acid resistance, was selected for *L. monocytogenes* RNA-seq analysis under acid stress.

### 3.3 Characterization of constitutive promoters via RNA-seq

The transcriptional profiling of all 2952 genes in wild-type EGD-e genome was performed via RNA-seq. Firstly, RNA quality, sequencing saturation, and redundant sequence distribution frequency were analyzed for quality control of transcriptome data (Fig. S1, Supporting Information). For two conditions of pH 7.4 and 5.5, each sample was sorted from highly expressed to low expressed according to the FPKM value. Compared with the normal condition of pH 7.4, 333 genes were up-regulated and 339 genes were down-regulated under the acid stress condition of pH 5.5 (Fig. 2a). The top 2.0 % of highly expressed genes under each condition were selected for gene overlap analysis. Among the 59 selected genes, 34 genes were co-expressed under two conditions (Fig. 2b). Since there are 15 genes collectively distributed under 6 different operons, 25 genes were chosen for cloning their promoters, which were listed in Table 1. And the promoter sequences were listed in Table S3.

**Table 1.** Selected 25 promoter regions from wild-type EGD-e via RNA-seq.

| Number | Gene | Length | CDS product | FPKM |  |
| --- | --- | --- | --- | --- | --- |
|  |  |  |  | pH 5.5 | pH 7.4 |
| 1 | lmo1634 | 501 | Aldehyde-alcohol dehydrogenase | 13392.83 | 52369.24 |
| 2 | lmo2653 | 108 | Elongation factor Tu | 41554.43 | 23758.48 |
| 3 | lmo2637 | 447 | Hypothetical protein | 5134.74 | 48137.68 |
| 4 | lmo2459 | 441 | Glyceraldehyde-3-phosphate dehydrogenase | 23215.88 | 18740.49 |
| 5 | lmo0045 | 278 | Single-strand binding protein | 21808.48 | 17971.34 |
| 6 | lmo1468 | 197 | Hypothetical protein | 17174.67 | 22337.2 |
| 7 | lmo2654 | 192 | Elongation factor G | 24542.36 | 10582.53 |
| 8 | lmo1439 | 300 | Superoxide dismutase | 27940.96 | 4269.49 |
| 9 | lmo2556 | 168 | Fructose-1,6-bisphosphate aldolase | 8363.89 | 17402.44 |
| 10 | lmo1541 | 156 | Hypothetical protein | 11821.15 | 11596.22 |
| 11 | lmo1364 | 198 | Cold-shock protein | 10301.34 | 7921.58 |
| 12 | lmo1003 | 306 | Phosphotransferase system enzyme I | 6824.64 | 10706.6 |
| 13 | lmo2612 | 495 | Preprotein translocase subunit | 10711.67 | 6707.29 |
| 14 | lmo2615 | 426 | 30S ribosomal protein S5 | 10020.13 | 5383.06 |
| 15 | lmo2610 | 386 | Translation initiation factor IF-1 | 9943.54 | 5379.53 |
| 16 | lmo2455 | 135 | Enolase | 6747.16 | 7920.58 |
| 17 | lmo2625 | 648 | 50S ribosomal protein L16 | 10782.57 | 3878.09 |
| 18 | lmo0248 | 126 | 50S ribosomal protein L11 | 6298.52 | 7050.71 |
| 19 | lmo1847 | 842 | Metal ABC transporter | 8066.96 | 4621.05 |
| 20 | lmo1787 | 207 | 50S ribosomal protein L19 | 6125.42 | 6160.1 |
| 21 | lmo2621 | 426 | 50S ribosomal protein L24 | 8505.77 | 3758.96 |
| 22 | lmo2016 | 288 | Cold-shock protein | 4096.5 | 8136.36 |
| 23 | lmo0210 | 293 | L-lactate dehydrogenase | 4121.04 | 7400.75 |
| 24 | lmo2411 | 1223 | Hypothetical protein | 6852.28 | 4470.22 |
| 25 | lmo0250 | 247 | 50S ribosomal protein L10 | 4045.77 | 5270.73 |

**Fig. 2.**
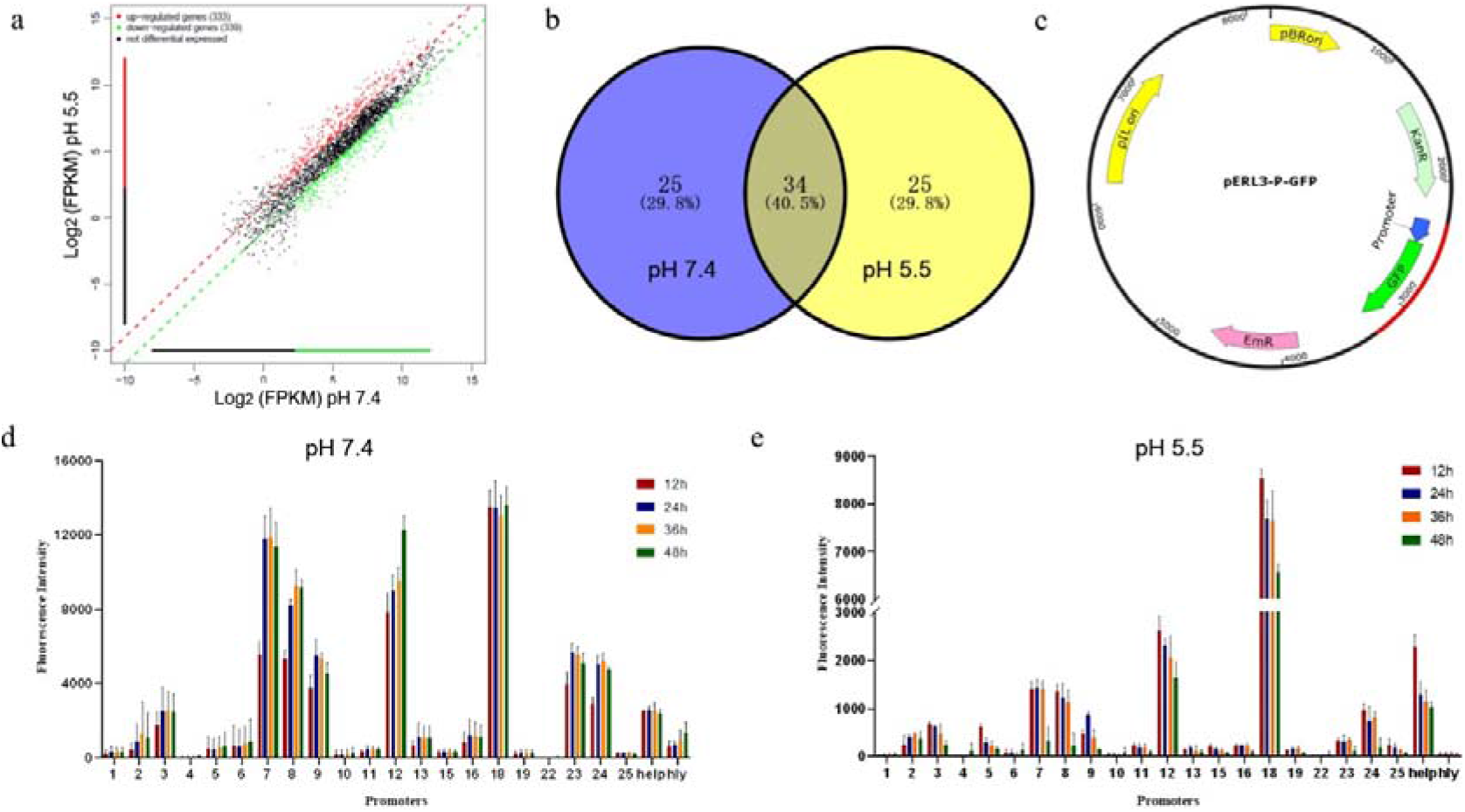
Characterization of constitutive promoters via RNA-seq. **a**. Scatter-plot of gene expression level at pH 5.5 and 7.4 via RNA-seq. The red spots indicate significantly up-regulated genes; the green spots indicate significantly down-regulated genes; black spots indicate nonsignificant genes. **b**. Venn diagram of the number of the top 2.0 % of the most highly expressed genes under two conditions by RNA-seq. **c**. Schematic diagram of the Promoter-GFP cassette for promoter screening. The sequence of Promoter-GFP cassette is highlighted in red color. Fluorescence intensity of GFP by different constitutive promoters in wild-type EGD-e at pH 7.4 (**d**) and 5.5 (**e**), respectively. The error bars indicate the standard deviations from three independent replicates.

To verify these putative constitutive promoters, the fluorescence intensities of GFP under different promoters in *L. monocytogenes* were measured for evaluating the promoter activity with constitutive promoters P_help_ and P_*hly*_ as references. According to the schematic diagram of plasmid construction (Fig. 2c), 21 fluorescent reporting plasmids with different promoters were successfully constructed and transformed into wild-type EGD-e. The strength of each promoter was similar at 12 h, 24 h, 36 h, and 48 h under the condition of pH 7.4 (Fig. 2d), confirming that they are constitutive promoters. Seven promoters P_7_, P_8_, P_9_, P_12_, P_18_, P_23_, and P_24_ showed high activities enhanced 1.8-fold to 5.4-fold compared with that of P_help_ in EGD-e. Under the condition of pH 5.5 (Fig. 2e), the fluorescence intensity of every well-characterized promoter has varying degrees of decline compared to that of pH 7.4. Nonetheless, the promoter P_18_ still showed a higher activity enhanced 5.3-fold compared to P_help_.

### 3.4 Fluorescent tracer of *L. monocytogenes* in vitro and in vivo

To assess the application potential of the well-characterized promoter, *L. monocytogenes* carrying fluorescent report plasmid with the strongest constitutive promoter P_18_ was selected for the tracer of bacteria in vitro and in vivo. The invasion of EGD-e and EGD-eΔ*actA/inlB* (pERL3-P_18_-GFP) in macrophage RAW264.7 could be observed (Fig. 3a). Further, the fluorescence signal of EGD-eΔ*actA/inlB* (pERL3-P_18_-GFP) was observed, and after dissection, could be assigned to livers (Fig. 3b), the major sites of listerial infections in mice. This indicates that the promoter P_18_ also has high activity in vivo.

**Fig. 3.**
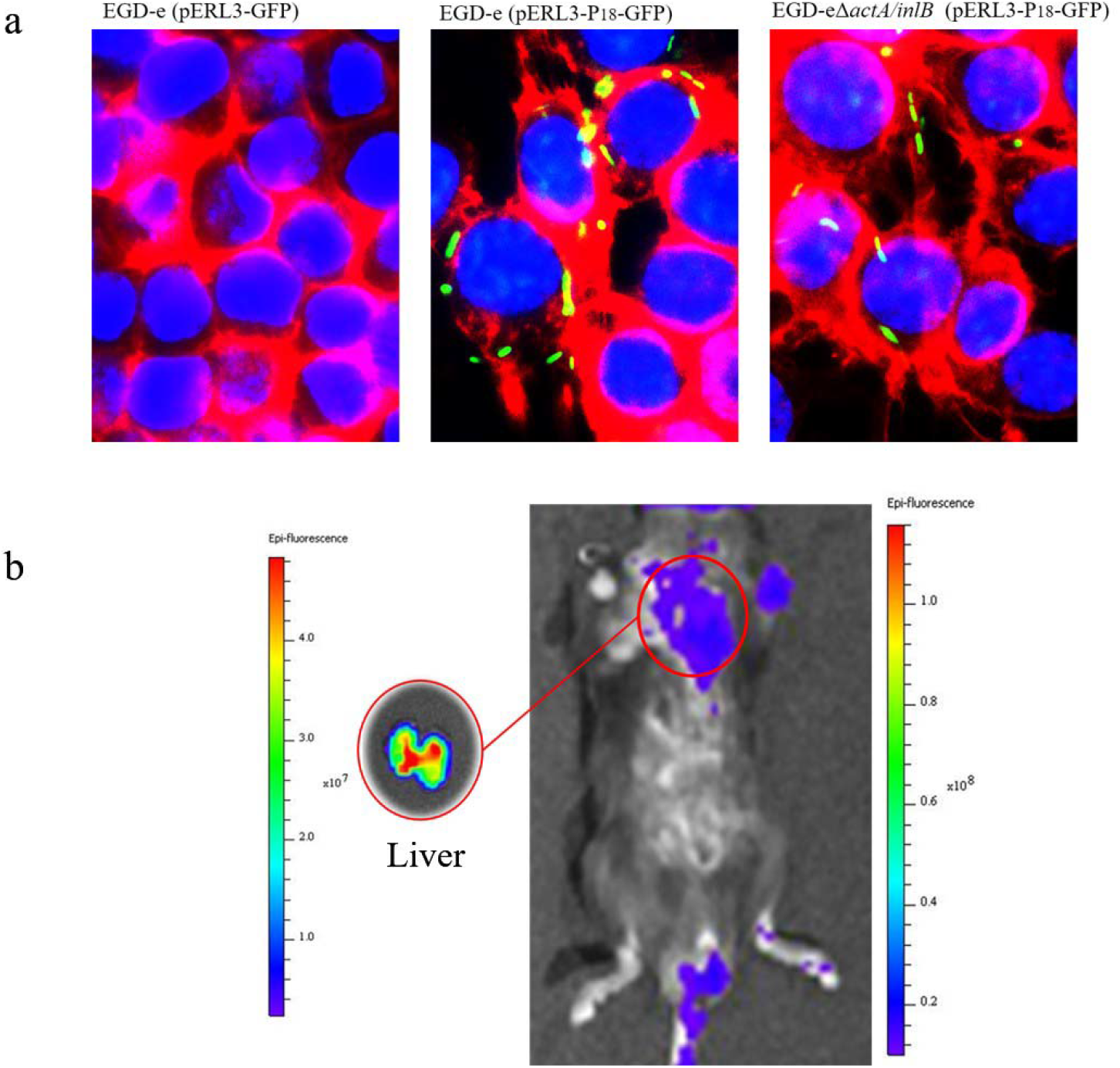
Fluorescent tracer of *L. monocytogenes* in vitro and in vivo. **a**. Fluorescent tracer of *L. monocytogenes* in macrophage RAW264.7. **b**. Fluorescent tracer of *L. monocytogenes* in vivo.

### 3.5 Measurement of UreB production

Attenuated *L. monocytogenes,* as a live vector for expressing foreign antigens in microbial immunotherapy research, needs a strong promoter to improve the expression of foreign antigens urgently. For this purpose, the promoters P_18_, P_7_, P_12_, P_9_, and P_24_ were selected to verify the expressing ability of the antigen UreB, a typical antigen of *H. pylori* with the promoters P_help_, P_*hly*_, and P_None_ (No promoter) as controls. As shown in Fig. 4a, the expression of UreB in EGD-eΔ*actA/inlB* under different promoters could be determined by anti-UreB polyclonal antibody using western blotting, which could be seen that the promoter P_18_ still had the highest ability to express UreB. With the gray-scale analysis of purified UreB, we carried out a quantitative analysis of UreB production under different promoters in EGD-eΔ*actA/inlB* (Fig. 4b). The promoters P_18_, P_7_, P_12_, P_9_, and P_24_ showed high production of UreB enhanced 1.1-fold to 8.3-fold compared with promoter P_help_. There was no doubt that the promoter P_18_ still performed noticeably well in the expression of antigen UreB.

**Fig. 4.**
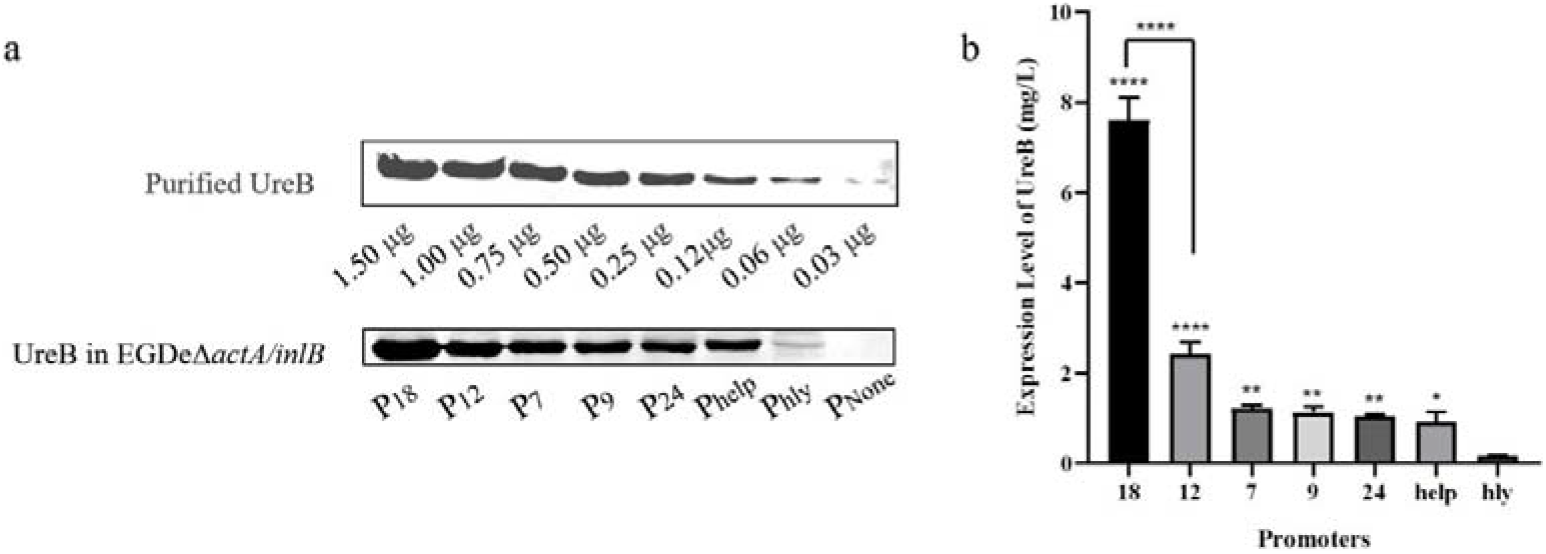
Measurement of UreB production. **a**. Measurement of UreB production in EGD-eΔ*actA/inlB* by Western Blotting. **b**. Quantitative determination of UreB production based on the gray scan. The error bars indicate the standard deviations from three independent replicates. Statistical significance was compared to the group of P_*hly*_: ns, no significant; *, P<0.05; **, P< 0.01; ****, P<0.0001.

## 4 Discussion

In metabolic engineering and synthetic biology, constitutive promoters play a great role in fine-tuning gene expression. The promoter with a proper strength could not only maximize the target production but also maintain the biological activity of the host engineering bacteria, which means that a mature engineering bacteria system needs a constitutive promoter library to select. Since *Listeria* is not a common engineered bacterium, the lack of available constitutive promoters in *Listeria* urges us to enrich this toolbox for *Listeria* species.

Considering that *L. monocytogenes*, as a vaccine vector, has to encounter the acid environment in the host, the promoter is required to maintain certain activity under acid stress conditions. Different acidity and processing time could result in different acid resistance of *L. monocytogenes*. In this work, the treatment of pH 5.5 for 3 h was selected as the acid stress condition of RNA-seq from all eight kinds of acid stress treatment. The bacteria under this treatment had the strongest acid resistance so that the down-regulated genes under acid stress could be filtered out as much as possible.

Based on RNA-seq and co-expression analysis under two conditions of pH 7.4 and pH 5.5, the characterizations of 21 promoters were identified by GFP reporter. Under the normal condition of pH 7.4, the activities of 7 promoters P_7_, P_8_, P_9_, P_12_, P_18_, P_23_, and P_24_ were higher than P_help_, a previously reported strong promoter. And under acid stress of pH 5.5, the fluorescence intensity of all promoters was decreased compared with that of pH 7.4, but there were still 2 promoters P_12_ and P_18,_ whose fluorescence intensity was higher than that of P_help_. As far as we know, this is the first well-characterized constitutive promoter library in *L. monocytogenes*.

Further, the well-characterized promoter P_18_ was evaluated in vitro and in vivo via *L. monocytogenes* strains carrying fluorescent reporter plasmid. Interestingly, the bacterial invasion in macrophage RAW264.7 and the bacterial enrichment in the liver of mice could be observed, which confirmed the feasibility of constitutive promoter P_18_ in the application. Also, the selected constitutive promoters P_18_, P_7_, P_12_, P_9_, and P_24_ with different strength had high activities in the production of the specific antigen UreB. Due to the dependence of antigen dose and antibody response, the higher antigen dose means the stronger immune response of specific antibodies[23]. So it was worth noting that the performance of constitutive promoter P_18_ was still far ahead in antigen production in *L. monocytogenes*.

In conclusion, we identified 21 candidate promoters from *L. monocytogenes* by RNA-seq under two conditions of pH 7.4 and pH 5.5. Based on the constructed fluorescent reporter system, 7 constitutive promoters showed great strength than that of P_help_, a previously reported strong promoter. To demonstrate their utility, 5 well-characterized constitutive promoters were used to successfully activate a foreign UreB biosynthetic pathway. In particular, the promoter activity in practical applications and the unprecedented antigen production of P_18_ were verified by the tracer of fluorescent reporter strains and UreB production. It could greatly enhance the effectiveness of live vector vaccines. Moreover, this study not only provides a useful toolkit for *Listeria* species but also offers a reference for constitutive promoter mining in other organisms.

## Authors’ contributions

Conceived and designed experiments: JF Ma, and Q Liu. Performed the experiments: JF Ma, and QY Ji. Analyzed the data: JF Ma, SY Wang, and JX Qiu. Wrote and revised the paper: JF Ma, and SY Wang.

## Acknowledgements

We thank Professor Qin Luo of Center China Normal University for the kind donation of *Listeria monocytogenes* EGD-e and plasmid pERL3.

## Declarations

### Ethical approval

This research does not contain any studies with human participants performed by any of the authors.

## Supplementary Information

**Fig. S1.**
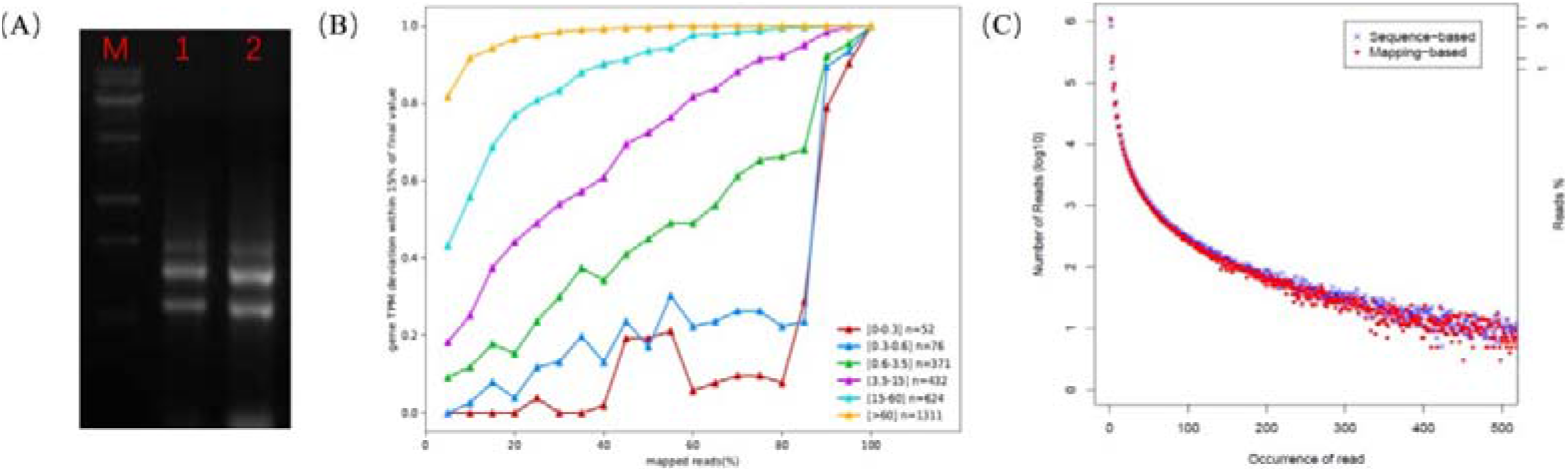
Descriptive statistics on RNA-seq. (A) Agarose gel electrophoresis of RNA samples from the conditions at pH 5.5 (1) and 7.4 (2). (B) Sequencing saturation analysis of RNA-seq. (C) Redundant sequence distribution frequency analysis of RNA-seq.

---

**Table S1.** Strains and plasmids in this study.

| Strains/plasmids | Description | Sources |
| --- | --- | --- |
| *E. coli* DH5α | Host for cloning | Purchased from Vazyme |
| EGD-e | *L. monocytogenes* wild-type | A gift from Qin Luo |
| EGD-eΔ*actA*/*inlB* | *L. monocytogenes* deleting gene *actA* and *inlB* | Our lab |
| Plasmids |  |  |
| pUC57-GFP | pUC57 derived, carrying a *gfp* gene, appropriate for *L. monocytogenes* after codon optimization | Our lab |
| pERL3 | *L. monocytogenes* shuttle vector, Em<sup>r</sup> | Our lab |
| pERL3-P<sub>1</sub>-GFP | pERL3 derived, carrying P<sub>1</sub> promoter-GFP cassette | This study |
| pERL3-P<sub>2</sub>-GFP | pERL3 derived, carrying P<sub>2</sub> promoter-GFP cassette | This study |
| pERL3-P<sub>3</sub>-GFP | pERL3 derived, carrying P<sub>3</sub> promoter-GFP cassette | This study |
| pERL3-P<sub>4</sub>-GFP | pERL3 derived, carrying P<sub>4</sub> promoter-GFP cassette | This study |
| pERL3-P<sub>5</sub>-GFP | pERL3 derived, carrying P<sub>5</sub> promoter-GFP cassette | This study |
| pERL3-P<sub>6</sub>-GFP | pERL3 derived, carrying P<sub>6</sub> promoter-GFP cassette | This study |
| pERL3-P<sub>7</sub>-GFP | pERL3 derived, carrying P<sub>7</sub> promoter-GFP cassette | This study |
| pERL3-P<sub>8</sub>-GFP | pERL3 derived, carrying P<sub>8</sub> promoter-GFP cassette | This study |
| pERL3-P<sub>9</sub>-GFP | pERL3 derived, carrying P<sub>9</sub> promoter-GFP cassette | This study |
| pERL3-P<sub>10</sub>-GFP | pERL3 derived, carrying P<sub>10</sub> promoter-GFP cassette | This study |
| pERL3-P<sub>11</sub>-GFP | pERL3 derived, carrying P<sub>11</sub> promoter-GFP cassette | This study |
| pERL3-P<sub>12</sub>-GFP | pERL3 derived, carrying P<sub>12</sub> promoter-GFP cassette | This study |
| pERL3-P<sub>13</sub>-GFP | pERL3 derived, carrying P<sub>13</sub> promoter-GFP cassette | This study |
| pERL3-P<sub>14</sub>-GFP | pERL3 derived, carrying P<sub>14</sub> promoter-GFP cassette | This study |
| pERL3-P<sub>15</sub>-GFP | pERL3 derived, carrying P<sub>15</sub> promoter-GFP cassette | This study |
| pERL3-P<sub>16</sub>-GFP | pERL3 derived, carrying P<sub>16</sub> promoter-GFP cassette | This study |
| pERL3-P<sub>17</sub>-GFP | pERL3 derived, carrying P<sub>17</sub> promoter-GFP cassette | This study |
| pERL3-P<sub>18</sub>-GFP | pERL3 derived, carrying P<sub>18</sub> promoter-GFP cassette | This study |
| pERL3-P<sub>19</sub>-GFP | pERL3 derived, carrying P<sub>19</sub> promoter-GFP cassette | This study |
| pERL3-P<sub>20</sub>-GFP | pERL3 derived, carrying P<sub>20</sub> promoter-GFP cassette | This study |
| pERL3-P<sub>21</sub>-GFP | pERL3 derived, carrying P<sub>21</sub> promoter-GFP cassette | This study |
| pERL3-P<sub>22</sub>-GFP | pERL3 derived, carrying P<sub>22</sub> promoter-GFP cassette | This study |
| pERL3-P<sub>23</sub>-GFP | pERL3 derived, carrying P<sub>23</sub> promoter-GFP cassette | This study |
| pERL3-P<sub>24</sub>-GFP | pERL3 derived, carrying P<sub>24</sub> promoter-GFP cassette | This study |
| pERL3-P<sub>25</sub>-GFP | pERL3 derived, carrying P<sub>25</sub> promoter-GFP cassette | This study |
| pERL3-P<sub>hly</sub>-GFP | pERL3 derived, carrying P<sub>hly</sub> promoter-GFP cassette | This study |
| pERL3-P<sub>help</sub>-GFP | pERL3 derived, carrying P<sub>help</sub> promoter-GFP cassette | This study |
| pERL3-P<sub>18</sub>-UreB | pERL3 derived, carrying P<sub>18</sub> promoter-UreB cassette | This study |
| pERL3-P<sub>7</sub>-UreB | pERL3 derived, carrying P<sub>7</sub> promoter-UreB cassette | This study |
| pERL3-P<sub>12</sub>-UreB | pERL3 derived, carrying P<sub>12</sub> promoter-UreB cassette | This study |
| pERL3-P<sub>9</sub>-UreB | pERL3 derived, carrying P<sub>9</sub> promoter-UreB cassette | This study |
| pERL3-P<sub>24</sub>-UreB | pERL3 derived, carrying P<sub>24</sub> promoter-UreB cassette | This study |
| pERL3-P<sub>hly</sub>-UreB | pERL3 derived, carrying P<sub>hly</sub> promoter-UreB cassette | This study |
| pERL3-P<sub>help</sub>-UreB | pERL3 derived, carrying P<sub>help</sub> promoter-UreB cassette | This study |

**Table S2.**
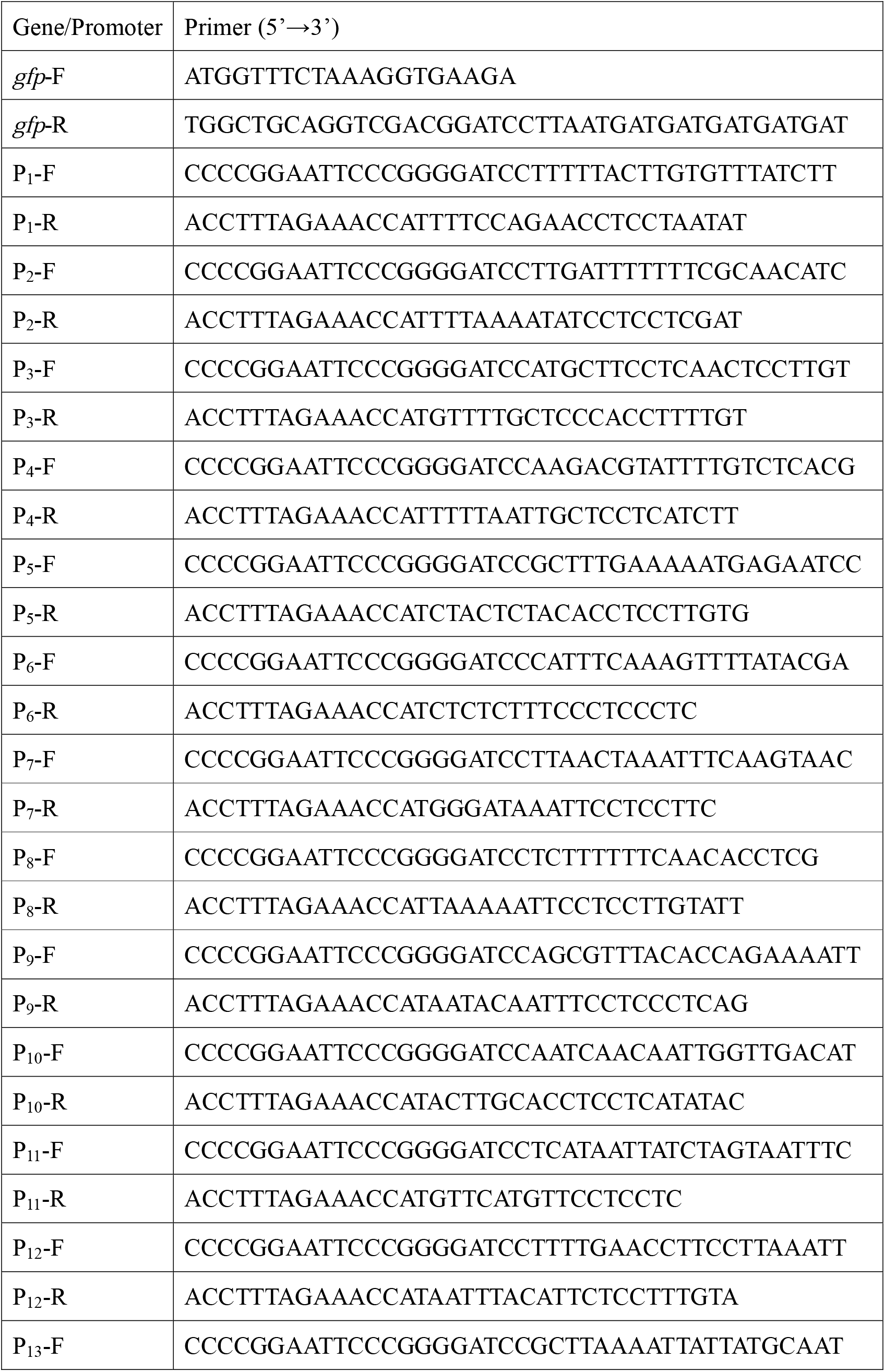
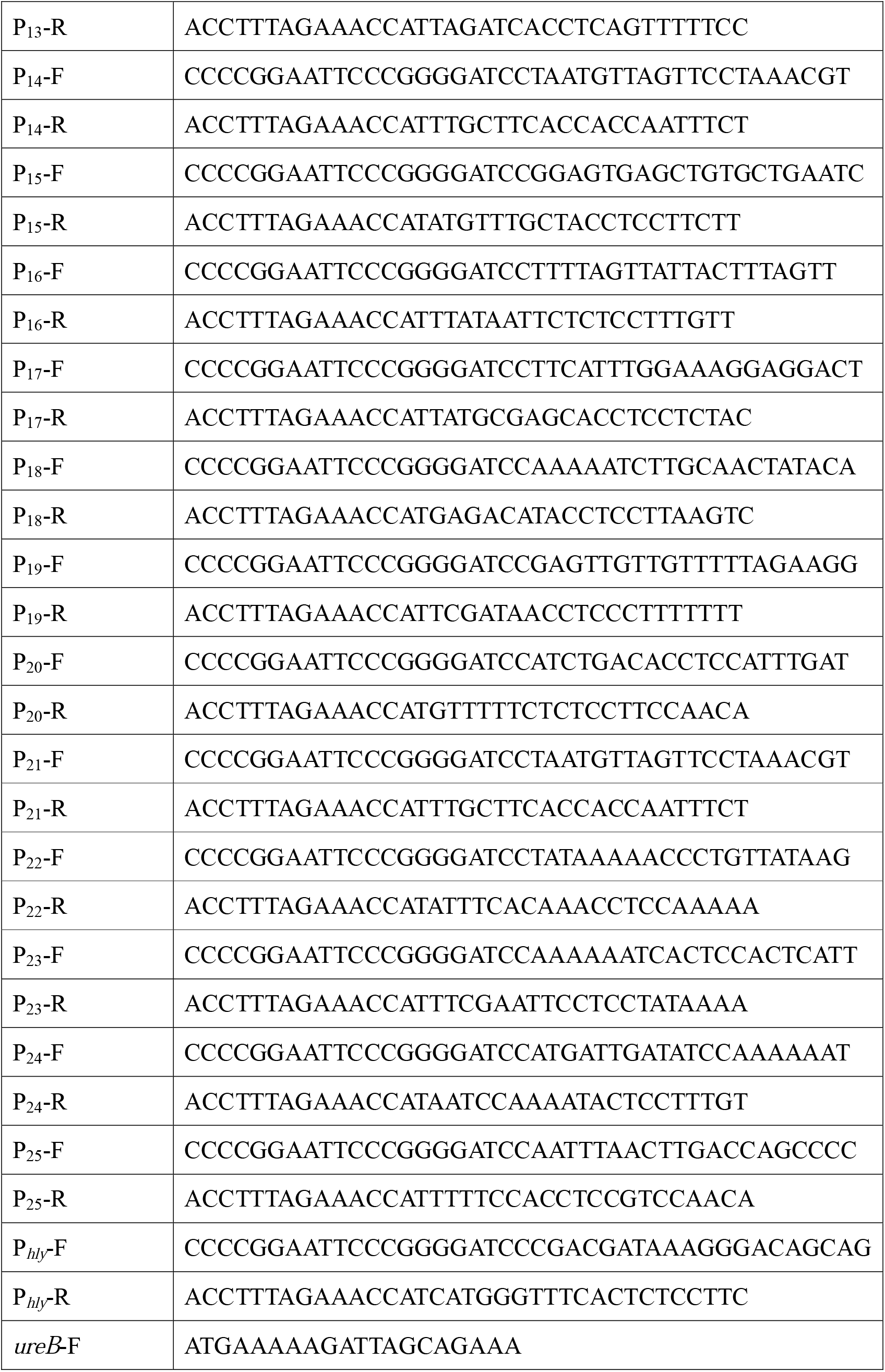
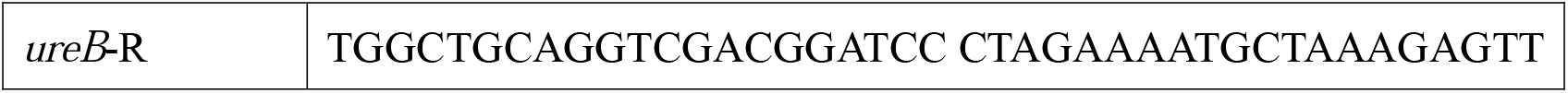
The primers used in this study.

**Table S3.**
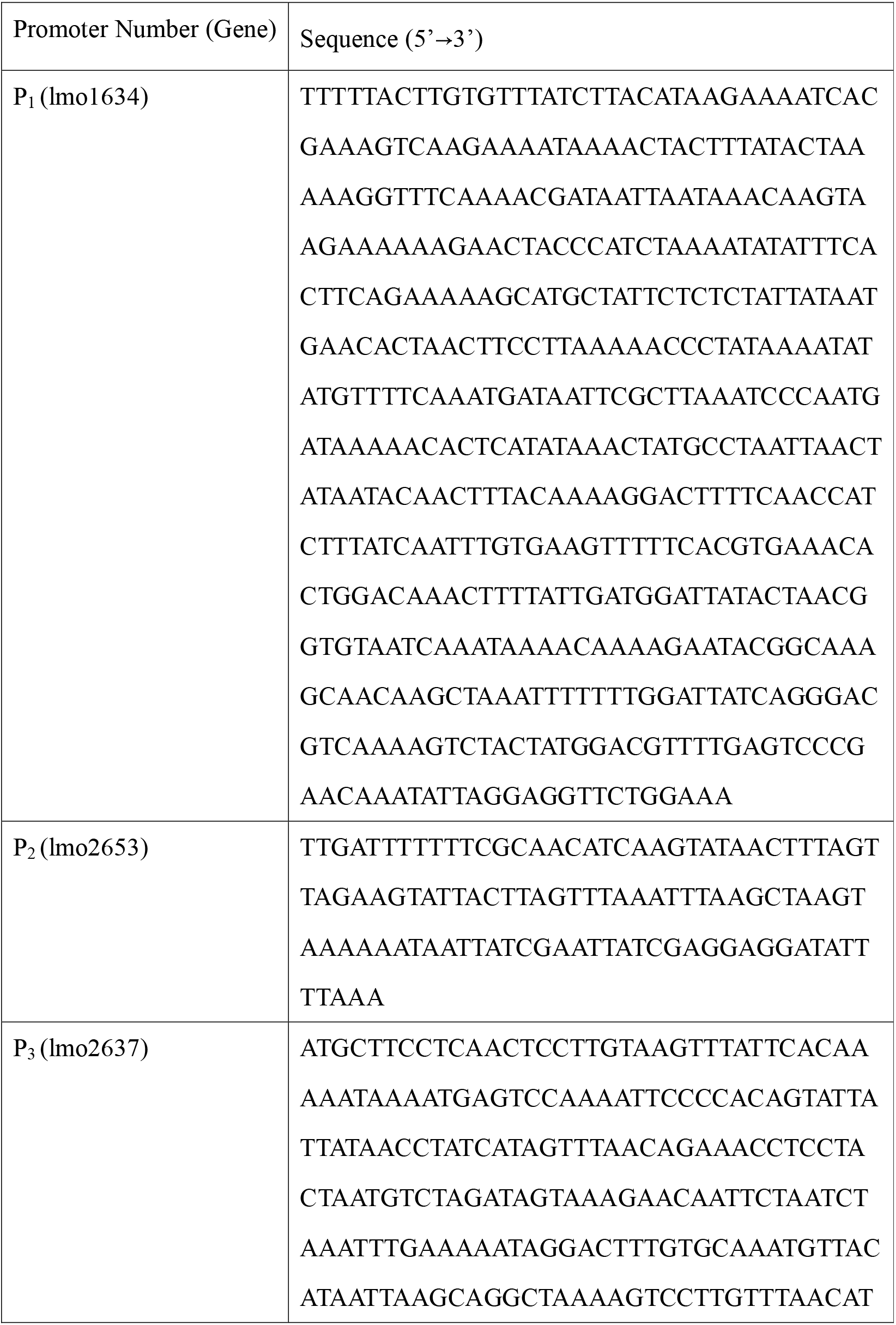
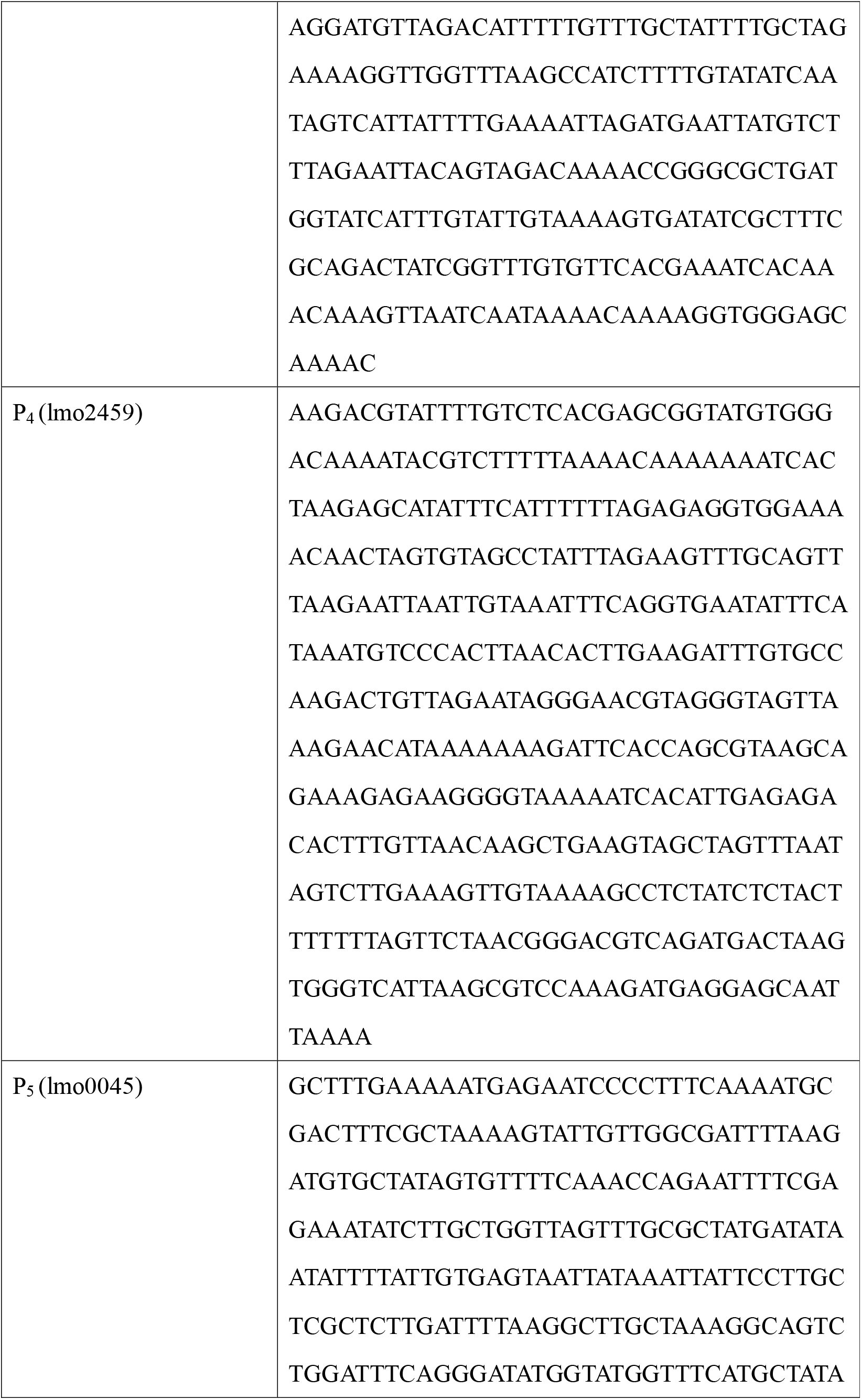
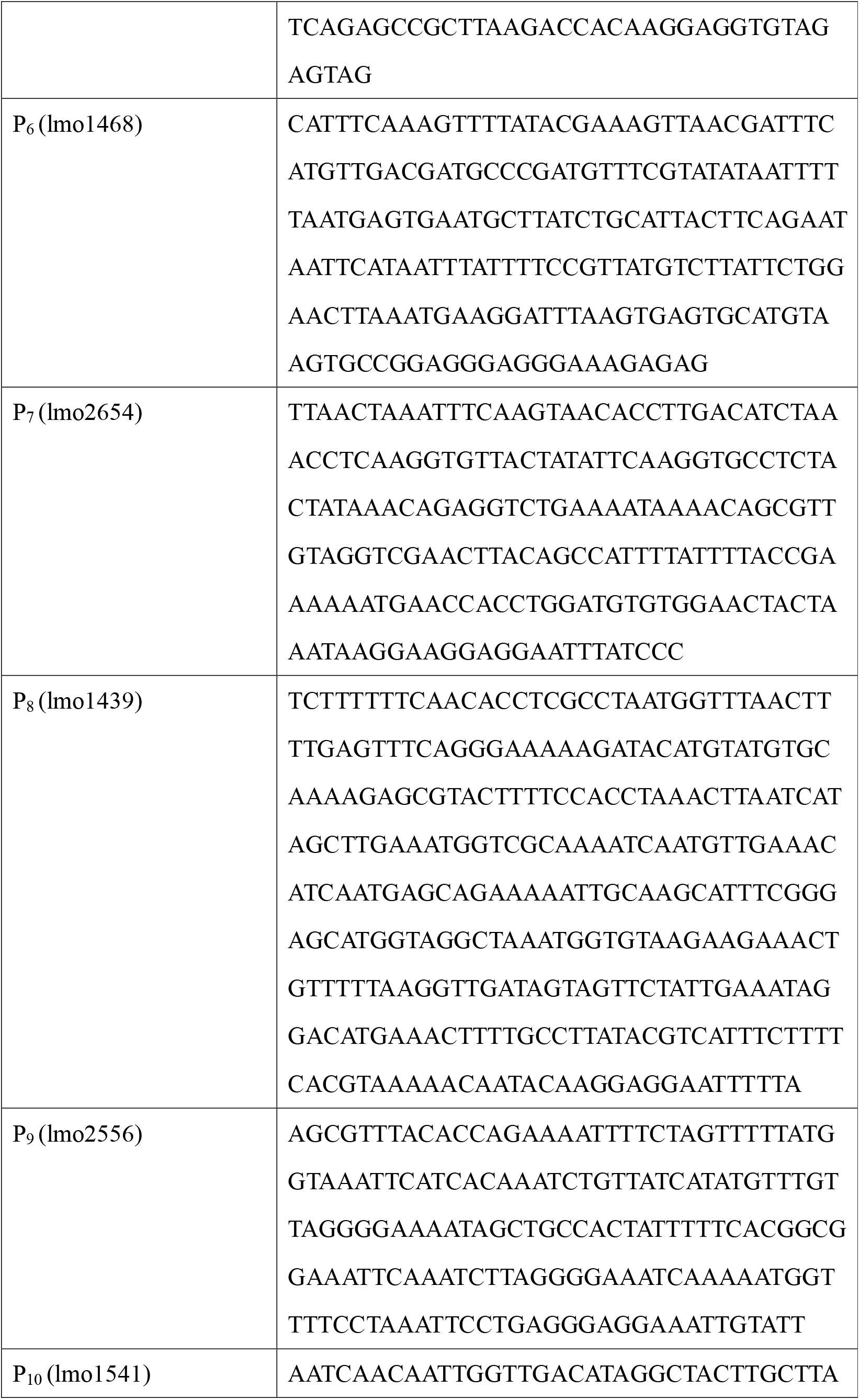
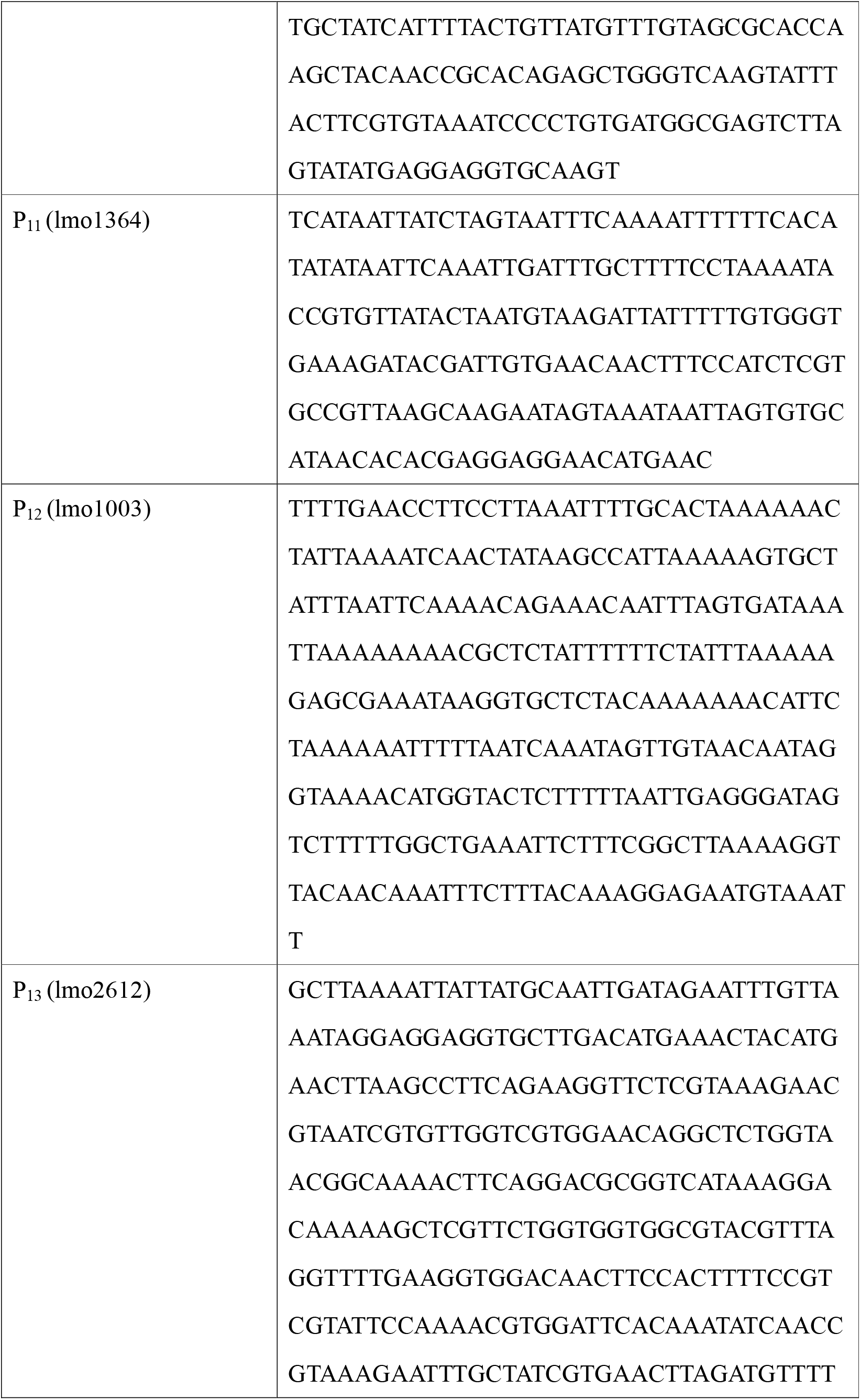
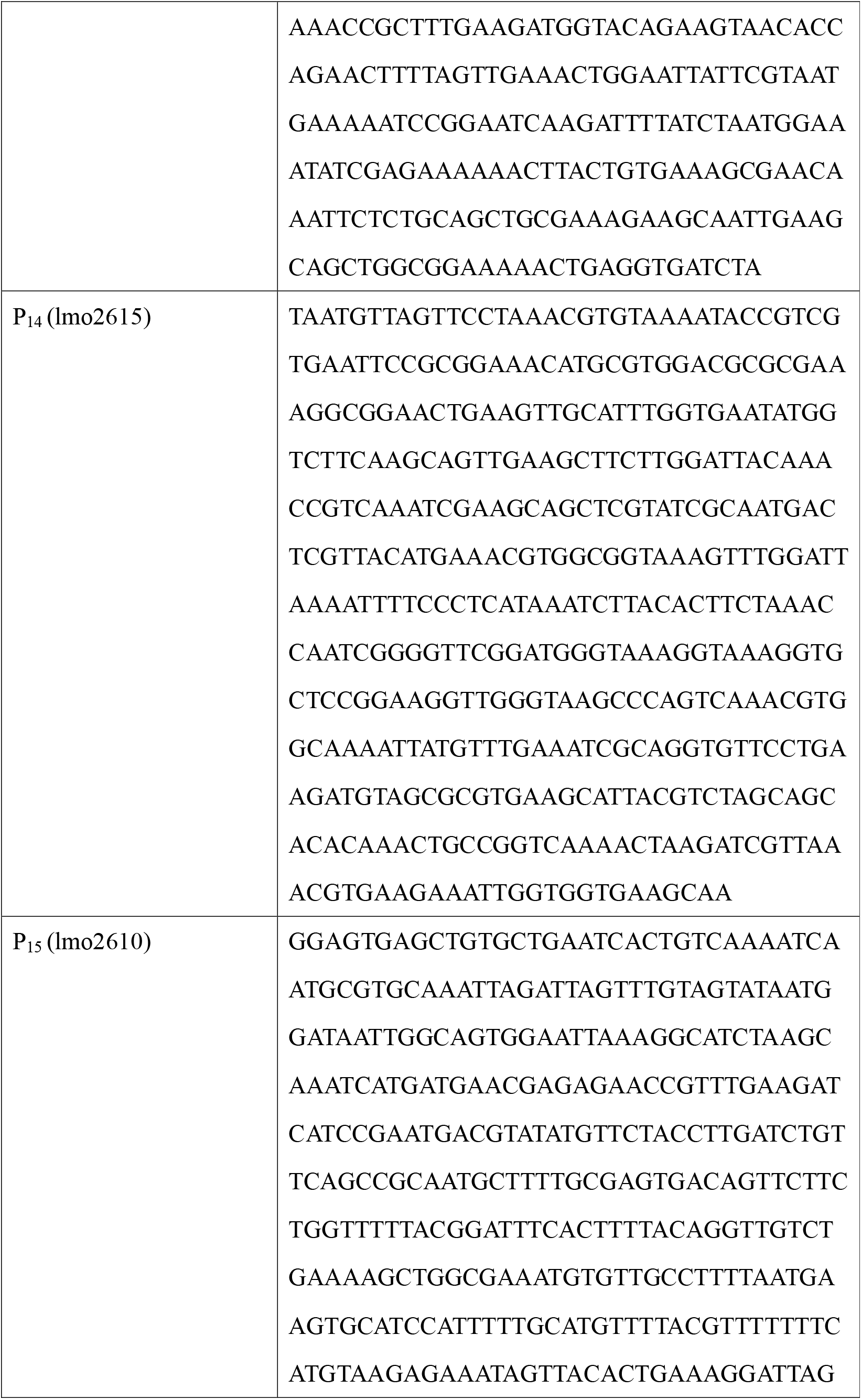
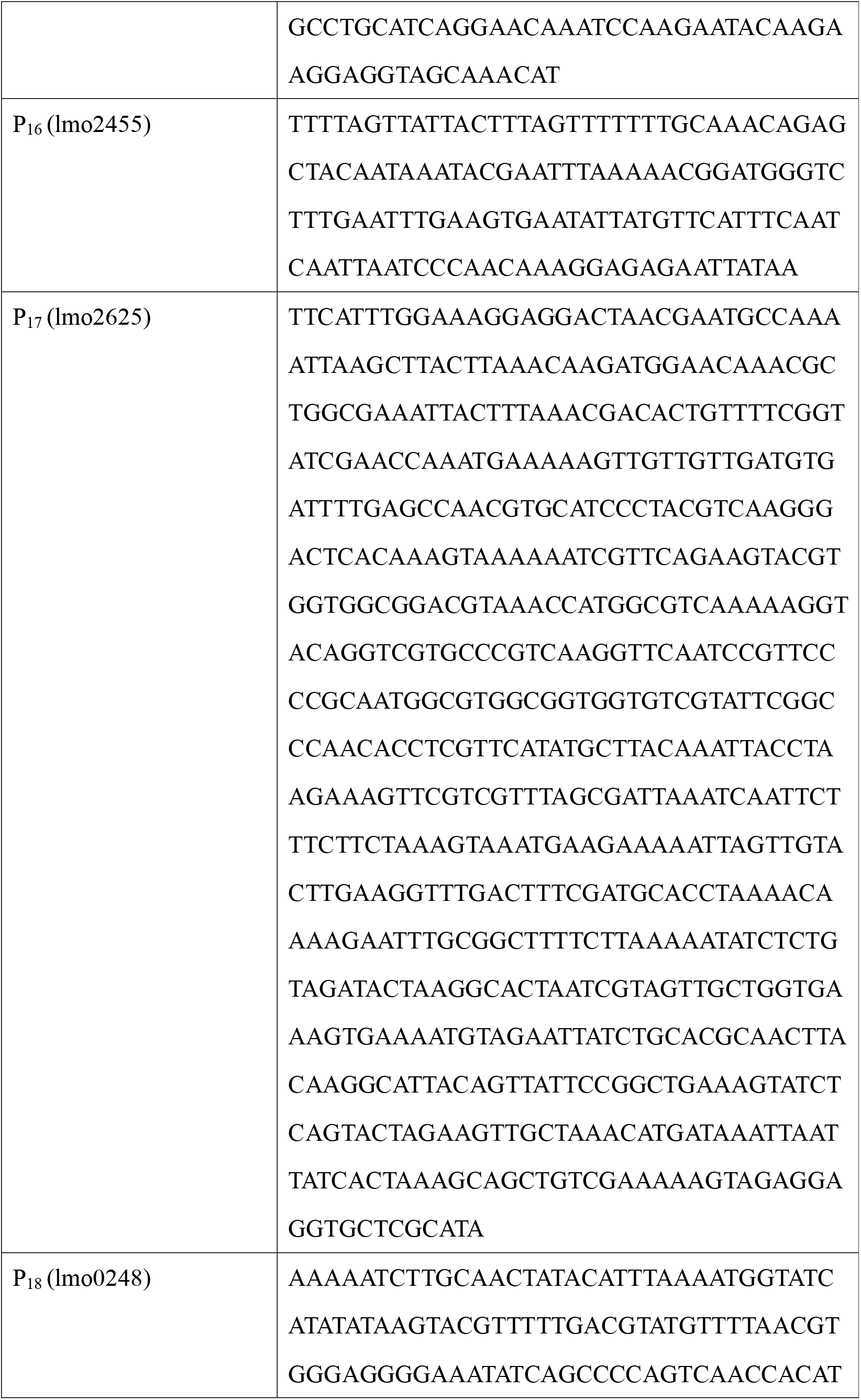
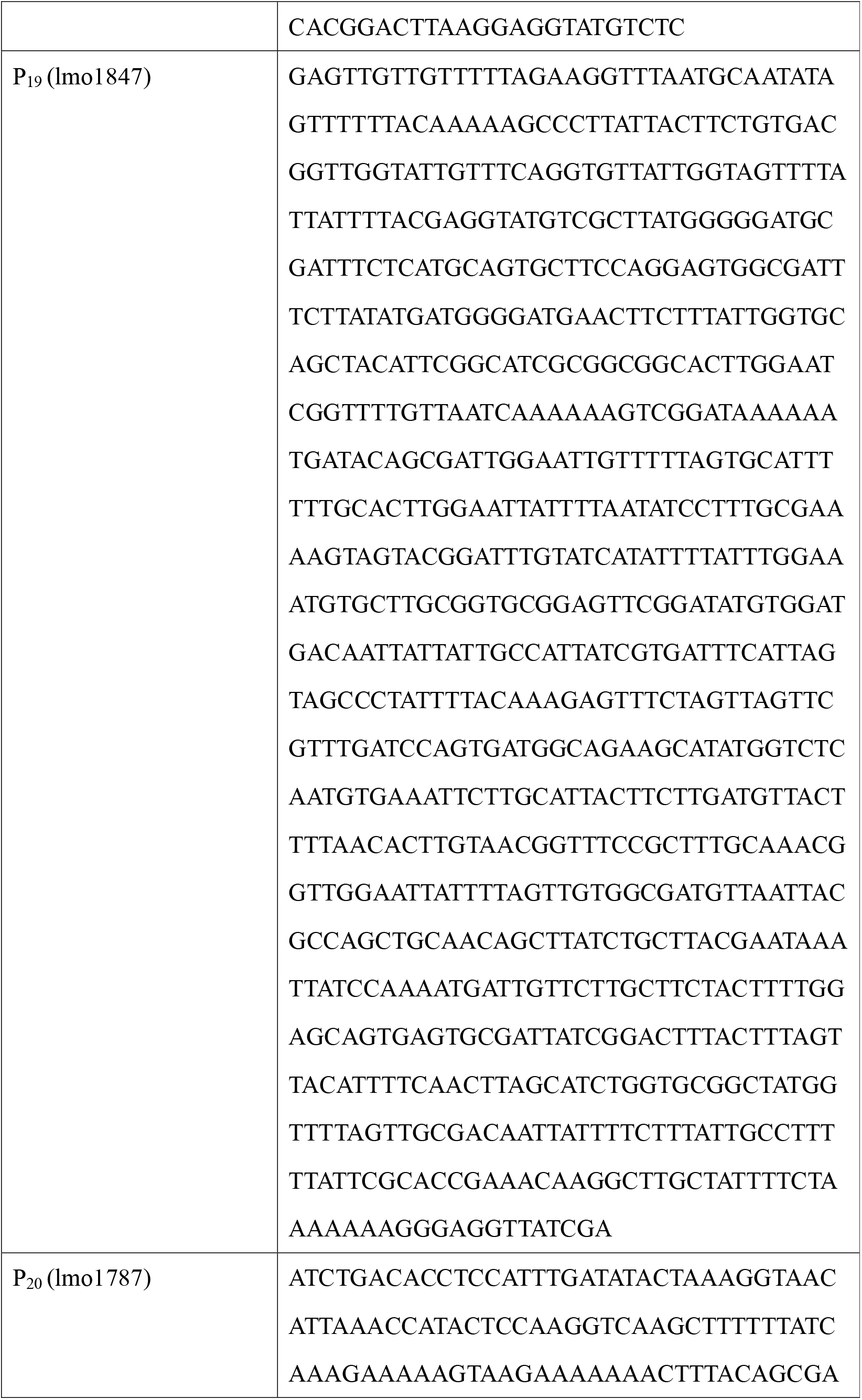
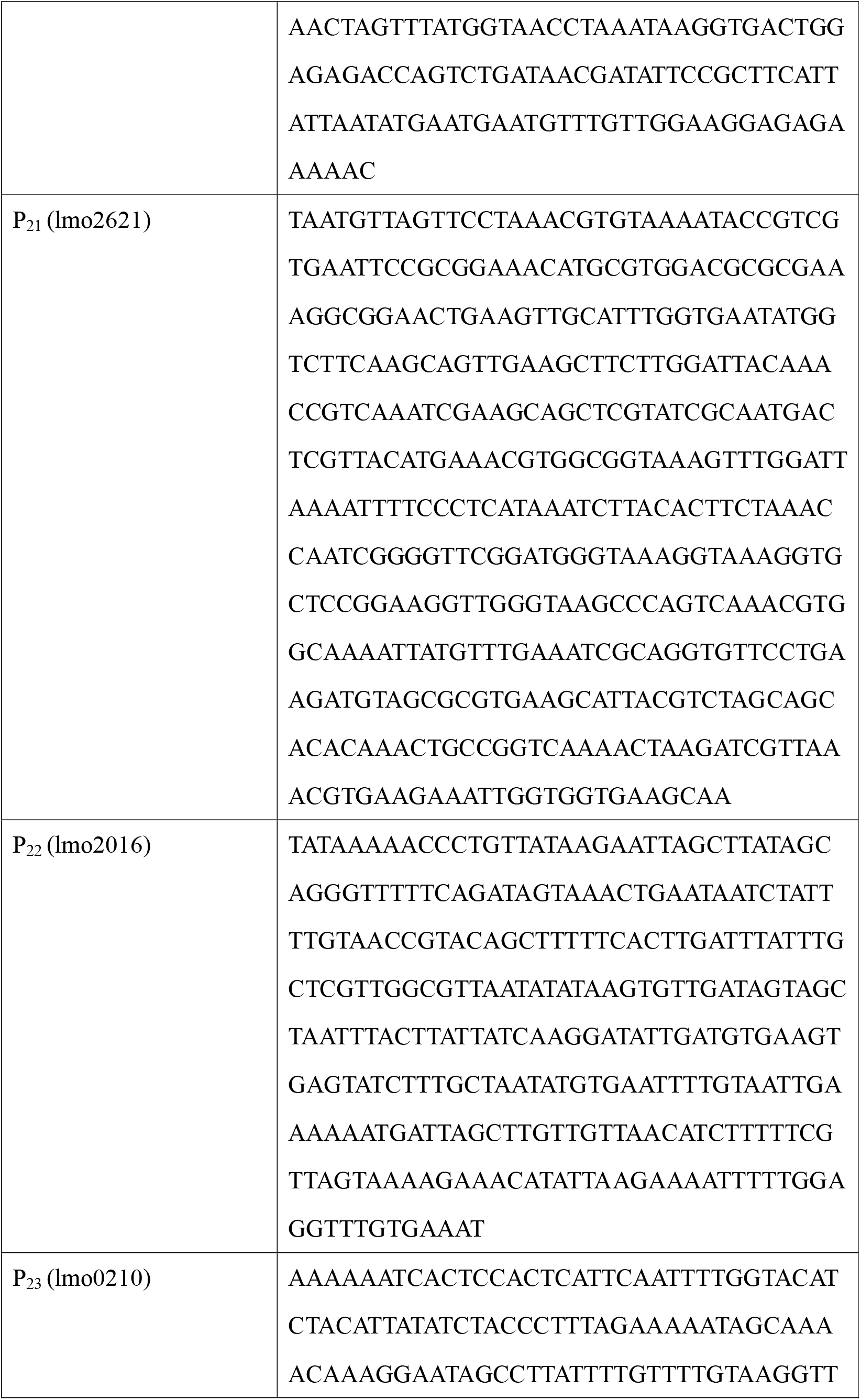
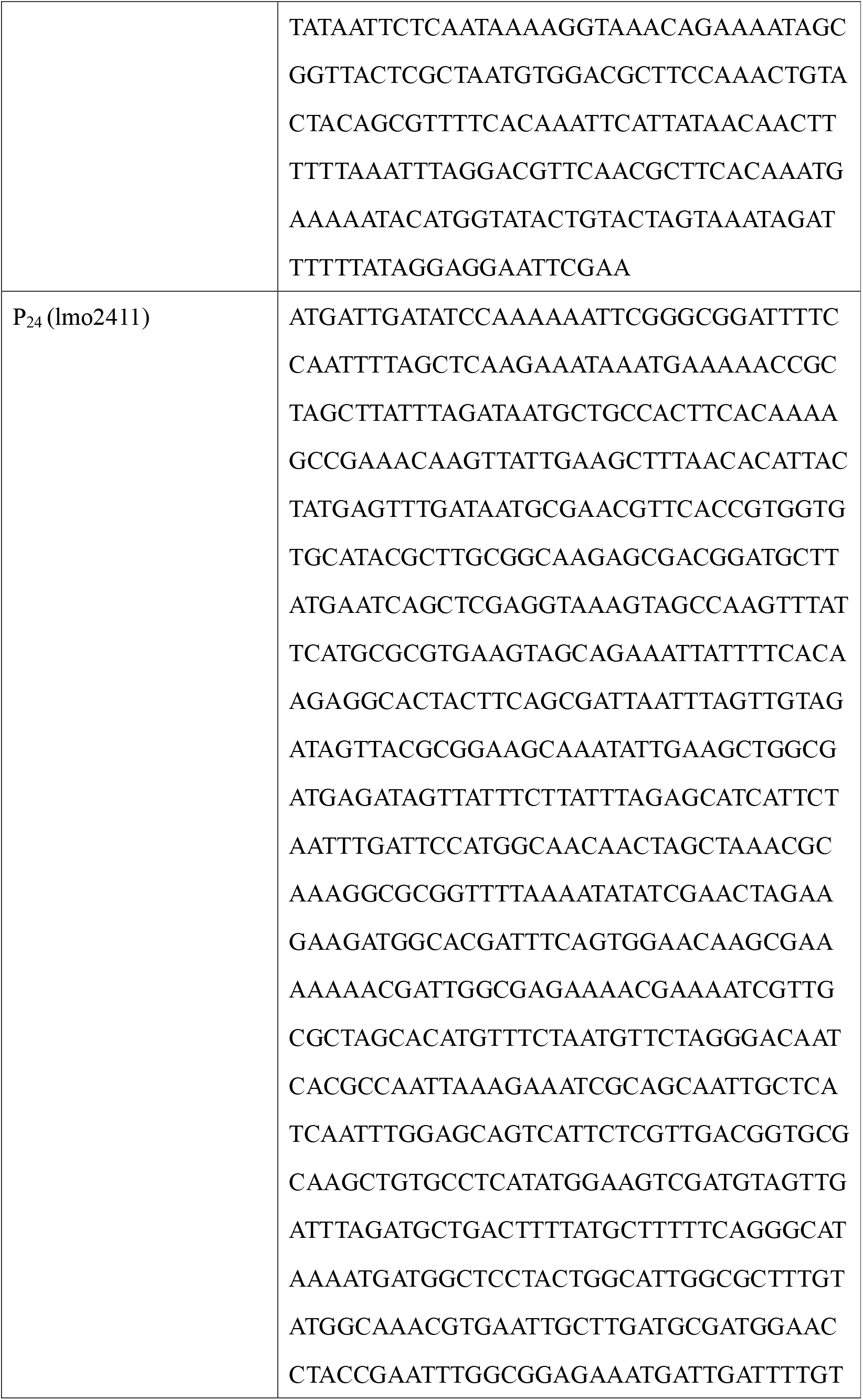
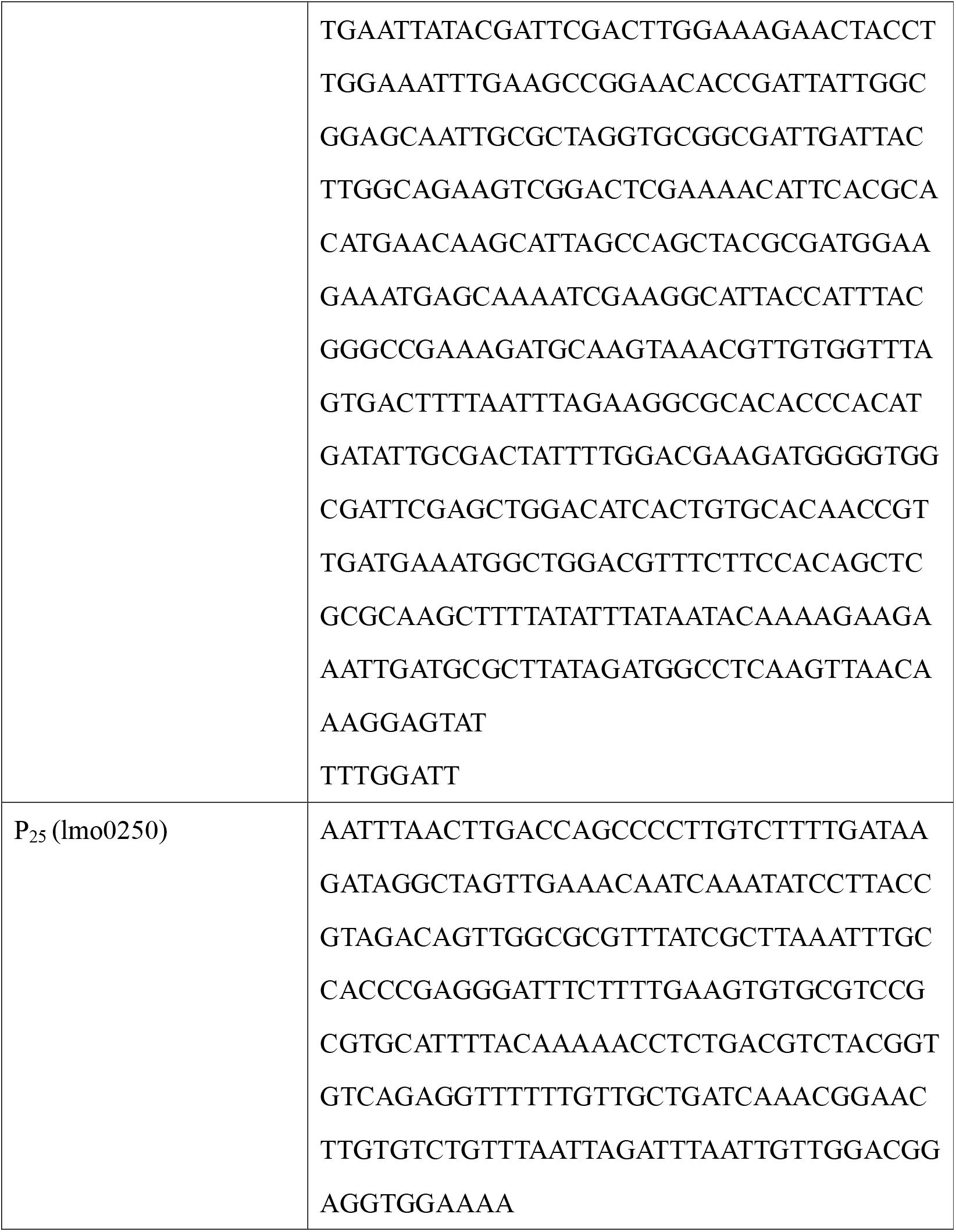
The sequences of selected promoters.

## References

[1] Guleria I, Pollard JW. Aberrant macrophage and neutrophil population dynamics and impaired Th1 response to *Listeria monocytogenes* in colony-stimulating factor 1-deficient mice. Infection and Immunity. 2001;69:1795–807.

[2] Mittrucker HW, Kohler A, Kaufmann SHE. Substantial in vivo proliferation of CD4(+) and CD8(+) T lymphocytes during secondary *Listeria monocytogenes* infection. European Journal of Immunology. 2000;30:1053–9.

[3] Stark FC, Sad S, Krishnan L. Intracellular Bacterial Vectors That Induce CD8(+) T Cells with Similar Cytolytic Abilities but Disparate Memory Phenotypes Provide Contrasting Tumor Protection. Cancer Research. 2009;69:4327–34.

[4] Yoshimura K, Laird LS, Chia CY, Meckel KF, Slansky JE, Thompson JM, et al. Live attenuated *Listeria monocytogenes* effectively treats hepatic colorectal cancer Metastases and is strongly enhanced by depletion of regulatory T cells. Cancer Research. 2007;67:10058–66.

[5] Le DT, Brockstedt DG, Nir-Paz R, Hampl J, Mathur S, Nemunaitis J, et al. A Live-Attenuated *Listeria* Vaccine (ANZ-100) and a Live-Attenuated Listeria Vaccine Expressing Mesothelin (CRS-207) for Advanced Cancers: Phase I Studies of Safety and Immune Induction. Clinical Cancer Research. 2012;18:858–68.

[6] Paterson Y, Johnson RS. Progress towards the use of *Listeria monocytogenes* as a live bacterial vaccine vector for the delivery of HIV antigens. Expert Rev Vaccines. 2004;3:S119–34.

[7] Chen Y, Yang D, Li S, Gao Y, Jiang R, Deng L, et al. Development of a *Listeria monocytogenes*-based vaccine against hepatocellular carcinoma (vol 31, pg 2140, 2012). Oncogene. 2012;31:4810-.

[8] Yang Y, Hou J, Lin Z, Zhuo H, Chen D, Zhang X, et al. Attenuated *Listeria monocytogenes* as a cancer vaccine vector for the delivery of CD24, a biomarker for hepatic cancer stem cells. Cellular & Molecular Immunology. 2014;11:184–96.

[9] Ding C, Liu Q, Li J, Ma J, Wang S, Dong Q, et al. Attenuated *Listeria monocytogenes* protecting zebrafish (Danio rerio) against Vibrio species challenge. Microbial Pathogenesis. 2019;132:38–44.

[10] Riedel CU, Monk IR, Casey PG, Morrissey D, O'Sullivan GC, Tangney M, et al. Improved luciferase tagging system for Listeria monocytogenes allows real-time monitoring in vivo and in vitro. Applied and Environmental Microbiology. 2007;73:3091–4.

[11] Huang QY, Roberts M. Nonspecific Phospholipase C Activity from *Listeria Monocytogenes* is Activated by Phagosome Acidification. Faseb J. 2013;27:1.

[12] Gallagher FA, Kettunen MI, Day SE, Hu DE, Ardenkjaer-Larsen JH, in't Zandt R, et al. Magnetic resonance imaging of pH in vivo using hyperpolarized C-13-labelled bicarbonate. Nature. 2008;453:940–U73.

[13] Spears PA, Suyemoto MM, Hamrick TS, Wolf RL, Havell EA, Orndorff PE. In Vitro Properties of a *Listeria monocytogenes* Bacteriophage-Resistant Mutant Predict Its Efficacy as a Live Oral Vaccine Strain. Infection and Immunity. 2011;79:5001–9.

[14] Sun J, Shao Z, Zhao H, Nair N, Wen F, Xu J-H, et al. Cloning and characterization of a panel of constitutive promoters for applications in pathway engineering in *Saccharomyces cerevisiae*. Biotechnology and Bioengineering. 2012;109:2082–92.

[15] Li S, Wang J, Li X, Yin S, Wang W, Yang K. Genome-wide identification and evaluation of constitutive promoters in *Streptomycetes*. Microbial Cell Factories. 2015;14.

[16] Partow S, Siewers V, Bjorn S, Nielsen J, Maury J. Characterization of different promoters for designing a new expression vector in *Saccharomyces cerevisiae*. Yeast. 2010;27:955–64.

[17] Luo Y, Zhang L, Barton KW, Zhao H. Systematic Identification of a Panel of Strong Constitutive Promoters from *Streptomyces albus*. Acs Synthetic Biology. 2015;4:1001–10.

[18] Kong L-H, Xiong Z-Q, Song X, Xia Y-J, Zhang N, Ai L-Z. Characterization of a Panel of Strong Constitutive Promoters from *Streptococcus thermophilus* for Fine-Tuning Gene Expression. Acs Synthetic Biology. 2019;8:1469–72.

[19] Jin Z-J, Zhou L, Sun S, Cui Y, Song K, Zhang X, et al. Identification of a Strong Quorum Sensing- and Thermo-Regulated Promoter for the Biosynthesis of a New Metabolite Pesticide Phenazine-1-carboxamide in *Pseudomonas* strain PA1201. ACS synthetic biology. 2020;9:1802–12.

[20] Trapnell C, Williams BA, Pertea G, Mortazavi A, Kwan G, van Baren MJ, et al. Transcript assembly and quantification by RNA-Seq reveals unannotated transcripts and isoform switching during cell differentiation. Nature Biotechnology. 2010;28:511–U174.

[21] Stack HM, Gahan CGM, Hill C. A novel promoter trap identifies *Listeria monocytogenes* promoters expressed at a low pH within the macrophage phagosome. Fems Microbiology Letters. 2007;274:139–47.

[22] Cheng C, Yang Y, Dong Z, Wang X, Fang C, Yang M, et al. *Listeria monocytogenes* varies among strains to maintain intracellular pH homeostasis under stresses by different acids as analyzed by a high-throughput microplate-based fluorometry. Frontiers in Microbiology. 2015;6.

[23] Li H, Limenitakis JP, Greiff V, Yilmaz B, Scharen O, Urbaniak C, et al. Mucosal or systemic microbiota exposures shape the B cell repertoire. Nature. 2020;584:274–+.

